# Genomic Study of Taste Perception Genes in African Americans Reveals SNPs Linked to Alzheimer’s Disease

**DOI:** 10.1101/2024.08.10.607452

**Authors:** Paule Valery Joseph, Malak Abbas, Gabriel Goodney, Ana Diallo, Amadou Gaye

**Affiliations:** National Institute on Alcohol Abuse and Alcoholism, National Institue of Nursing Research, Sensory Science and Metabolism Unit, Biobehavioral Branch, National Institutes of Health, Bethesda, MD, USA; National Human Genome Research Institute, National Institutes of Health, Bethesda, MD, USA; Department of Pharmacotherapy & Outcomes Science, Virginia Commonwealth University, Richmond, VA

**Keywords:** Taste Genes, GWAS, Transcriptome, Proteome, Alzheimer’s Disease, African American

## Abstract

**Background:** While previous research has shown the potential links between taste perception pathways and brain-related conditions, the area involving Alzheimer’s disease remains incompletely understood. Taste perception involves neurotransmitter signaling, including serotonin, glutamate, and dopamine. Disruptions in these pathways are implicated in neurodegenerative diseases. The integration of olfactory and taste signals in flavor perception may impact brain health, evident in olfactory dysfunction as an early symptom in neurodegenerative conditions. Shared immune response and inflammatory pathways may contribute to the association between altered taste perception and conditions like neurodegeneration, present in Alzheimer’s disease.

**Methods:** This study consists of an exploration of expression-quantitative trait loci (eQTL), utilizing whole-blood transcriptome profiles, of 28 taste perception genes, from a combined cohort of 475 African American subjects. This comprehensive dataset was subsequently intersected with single-nucleotide polymorphisms (SNPs) identified in Genome-Wide Association Studies (GWAS) of Alzheimer’s Disease (AD). Finally, the investigation delved into assessing the association between eQTLs reported in GWAS of AD and the profiles of 741 proteins from the Olink Neurological Panel.

**Results:** The eQTL analysis unveiled 3,547 statistically significant SNP-Gene associations, involving 412 distinct SNPs that spanned all 28 taste genes. In 17 GWAS studies encompassing various traits, a total of 14 SNPs associated with 12 genes were identified, with three SNPs consistently linked to Alzheimer’s disease across four GWAS studies. All three SNPs demonstrated significant associations with the down-regulation of *TAS2R41*, and two of them were additionally associated with the down-regulation of *TAS2R60*. In the subsequent pQTL analysis, two of the SNPs linked to *TAS2R41* and *TAS2R60* genes (rs117771145 and rs10228407) were correlated with the upregulation of two proteins, namely EPHB6 and ADGRB3.

**Conclusions:** Our investigation introduces a new perspective to the understanding of Alzheimer’s disease, emphasizing the significance of bitter taste receptor genes in its pathogenesis. These discoveries set the stage for subsequent research to delve into these receptors as promising avenues for both intervention and diagnosis. Nevertheless, the translation of these genetic insights into clinical practice requires a more profound understanding of the implicated pathways and their pertinence to the disease’s progression across diverse populations.

## INTRODUCTION

Taste perception is a complex sensory phenomenon that involves the detection and interpretation of various chemical stimuli by specialized receptors located on the tongue and oral cavity ^1^. The human sense of taste, or gustation, is primarily categorized into five modalities: sweet, salty, sour, bitter, and umami. The intricate interplay between taste perception and the brain constitutes a process, integrating peripheral sensory signals with central neural processing. Neuroimaging studies, such as functional magnetic resonance imaging and positron emission tomography, have provided valuable insights into the neural mechanisms underlying taste perception ^2,3^. These studies reveal the involvement of various brain regions in different aspects of taste processing, highlighting the distributed nature of the taste neural network.

There is evidence to suggest potential links between taste perception pathways and brain-related conditions ^4,5^; however, the connections are still not fully understood. For instance, taste perception involves neurotransmitter signaling in the gustatory system, particularly through molecules such as serotonin, glutamate, and dopamine ^6^. Disruptions in these neurotransmitter systems have been implicated in mood disorders, schizophrenia, and neurodegenerative diseases ^7,8^. The integration of olfactory and taste signals in flavor perception may have implications for brain health. For example, olfactory dysfunction is a common early symptom in neurodegenerative conditions like Alzheimer’s and Parkinson’s diseases ^9^. Futhermore, inflammation plays a role in both alterations in taste perception and neuroinflammatory conditions. Shared pathways involving immune response and inflammatory mediators may contribute to the link between altered taste perception and conditions like depression and neurodegeneration ^10,11^.

Alzheimer’s disease (AD) is a progressive neurodegenerative disorder that significantly impacts cognitive function and daily living activities. As the most common cause of dementia, AD is characterized by the accumulation of amyloid-beta plaques and tau tangles, leading to neuronal loss and a decline in cognitive abilities ^12^. Alongside these well-documented features, AD is increasingly associated with sensory dysfunctions, including alterations in taste and smell perception ^9,13^. These sensory changes are not just secondary symptoms but are thought to be integral to the disease process, potentially serving as early indicators of neurodegeneration ^9^.

Taste alterations in individuals with Alzheimer’s diseases have been documented in various studies ^13-15^. Individuals with AD often experience changes in food preferences, decreased taste sensitivity or increase taste preference for sweet and salty tastes, and sometimes a complete loss of taste perception. These alterations can lead to nutritional imbalances and affect the quality of life and overall health ^16^.

The underlying mechanisms are believed to involve both central and peripheral pathways, including changes in taste bud integrity, central processing of taste information and taste receptor expression.

Genetic factors influencing taste receptor expression may have broader implications for many diseases including brain conditions ^17-20^. Polymorphisms in taste receptor genes loci may contribute to variations in gene expression and protein levels that could be associated with neurodenerative conditions such as Alzheimer’s disease (AD) ^21^. Functional genomics approaches, leveraging transcriptomics and proteomics can serve as valuable supplements to genetic inquiries and offer perspectives into the molecular mechanisms that link polymorphisms associated with taste perception to brain diseases. Such strategies can help decipher the complex biological pathways linked to the initiation and advancement of those diseases.

Notably, AD and associated sensory dysfunctions may vary significantly across different populations, influenced by a complex interplay of genetic, environmental, and social factors. African Americans are disproportionately affected by AD, experiencing higher incidence rates and more severe cognitive deterioration compared to other racial groups^22,23^. These disparities are not fully understood and are attributed to factors including genetic predisposition, health comorbidities, socioeconomic status, and access to healthcare. Understanding the specific genetic and molecular basis of taste alterations in an African American cohort can provide insights into the unique progression of AD in this population and highlight potential areas for targeted intervention and care. The objectives of this project are twofold: (1) identify expression-quantitative trait loci (eQTL) impacting the expression of taste-related genes that have been reported in genome-wide association studies (GWAS) of brain conditions, and (2) evaluate the connections between these variants and proteins involved in neurological processes relevant to neurodegenerative conditions and specifically AD.

## MATERIAL AND METHODS

### Phenotype data

The GENomics, Environmental FactORs and the Social DEterminants of Cardiovascular Disease in African-Americans STudy (GENE-FORECAST) is a research platform that establishes a strategic, multi-omics systems biology approach amenable to the deep, multi-dimensional characterization of minority health and disease in AA (African American). GENE-FORECAST is designed to create a cohort based on a community-based sampling frame of U.S.-born, AA men and women (ages 21-65) recruited from the metropolitan Washington D.C. area.

The Minority Health Genomics and Translational Research Bio-Repository Database (MH-GRID) project is a study of hypertension (HTN) in AA aged 30 to 55 years. The data included in this analysis is from an MH-GRID sub-study of samples from the Morehouse School of Medicine (MSM), in Atlanta (Georgia).

### Transcriptome data

The transcriptome data consisted of the mRNA sequencing data of whole blood (buffy coat). Total RNA extraction was carried out using MagMAXTM for Stabilized Blood Tubes RNA Isolation Kit as recommended by vendor (Life Technologies, Carlsbad, CA). For library preparation, total RNA samples are concentration normalized, and ribosomal RNA (rRNA) is removed. Illumina paired end sequencing was performed on HiSeq2000 analyzer (Illumina, USA) with an average sequencing depth of 50 million reads per sample. The mRNA expression was quantified using a bioinformatics pipeline developed by the Broad Institutes and used by the Genotype-Tissue Expression (GTEx). The pipeline is detailed in the GitHub software development platform ^24^. Transcripts that did not achieve an expression of one read count per million (CPM) in at least three samples were excluded. The expression data was normalized using the Trimmed Mean of M-values (TMM), an optimal method for read count data ^25^. Principal component analysis was conducted to identify and exclude sample and gene outliers. A set of 17,948 protein coding mRNA passed all quality control (QC) filters including a total of 28 taste perception related genes considered for this analysis. The 28 genes are listed in Supplemental Table T1A with their raw read counts reported the supplemental document named raw_count_data_for_RNA_expression and the distribution of the counts shown graphically in Supplemental Material M1. The genes consist of 22 TAS2R genes related to bitter taste perception, 3 SCNN1 genes related to salty taste perception and 3 TAS1R genes related to to umami (glutamate) perception.

### Genotype Data

For all the samples DNA was extracted from blood collected in PAXgene Blood DNA Tube and plated for genotyping on the Illumina Multi-Ethnic Genotyping Array (MEGA) version 2 which includes more than two million loci. The loci targeted by this Illumina product have been specified by the PAGE (Population Architecture in Genetics and Epidemiology) consortium and the ADPC (African Diaspora Power Chip) consortium. The raw intensity data were analyzed using Illumina proprietary software, the Illumina® GenomeStudio Genotyping Module™ Genotyping Module Software v2.0 and clustering for genotype assignment was of high quality (GenTrain Score > 0.9). Pre-analysis QC were carried out according to best practice. Briefly, markers with missing rate > 0.1 were exclude as well as those that failed Hardy-Weinberg Equilibrium test; principal component analysis (PCA) was conducted to identify and exclude sample outliers. Finally, samples with discordant sex information between genotype and phenotype data were also excluded. For the purpose of this work, unrelated samples that have proteome and/or transcriptome data in both cohorts were included in the analyses.

### Proteome data

The ethylenediamine tetraacetic acid (EDTA) plasma samples from the GENE-FORECAST and MH-GRID cohorts were sent to the Olink Proteomics Analysis Service in Boston, USA. Proteomic analyses were conducted collectively in a single batch, and the data were delivered in November 2022. The Explore 3072 assay, utilizing eight Explore 384 Olink panels (Cardiometabolic, Cardiometabolic II, Inflammation, Inflammation II, Neurology, Neurology II, Oncology, Oncology II), was run to assess the relative expression of a total of 2,947 proteins. Proximity Extension Assay (PEA) technology was conducted according to the Olink AB manufacturer procedures by the certified laboratory. Briefly, the technique relies on the use of antibodies labelled with unique DNA oligonucleotides that bind to their target protein present in the sample. The DNA oligonucleotides, when in proximity on a target protein, undergo hybridization and act as a template for DNA polymerase-dependent extension, forming a unique double-stranded DNA barcode proportionate to the initial protein concentration. Quantification of resulting DNA amplicons is accomplished through high-throughput DNA sequencing, generating a digital signal reflective of the number of DNA hybridization events corresponding to the protein concentration in the original sample. The measurement of protein levels is based on Normalized Protein eXpression (NPX) values, serving as a relative protein quantification unit. This quantification is normalized to account for systematic noise arising from sample processing and technical variation, leveraging internal controls and sample controls. NPX units are on a log2 scale, where a one NPX unit increase indicates a two-fold rise in the concentration of amplicons representing the target protein compared to the internal control. A total of 2,941 passed QC filtering and a subset of 741 of those from the neurological panel were included in the protein quantitative trait loci (pQTL) analysis with 585 samples (458 from GENE-FORECAST and 127 from MH-GRID) which have genotype and proteome data available.

### Statistical Analyses

#### eQTL analysis

The analysis included all bi-allelic single nucleotide polymorphisms (SNPs) with a minor allele frequency (MAF) ≥ 0.01 within the combined GENE-FROECAST and MH-GRID dataset (n = 475). Only SNPs within the cis-region (within one megabase) of the 28 taste perception genes in the transcriptome data were considered. The decision to focus on bi-allelic SNPs in the analyses is informed by various considerations, such as the superior accuracy and reliability associated with genotyping techniques for bi-allelic SNPs when compared to multi-allelic SNPs. This enhanced precision contributes to the robustness of the eQTL findings. Moreover, the emphasis on bi-allelic SNPs is strategically guided by their binary allele composition, which facilitates the interpretation of genetic effects and simplifies the identification and characterization of associations between specific alleles and mRNA and protein levels. A SNP was designated as an eQTL if the false-discovery rate (FDR) adjusted p-value of the association with a gene was ≤ 0.05.

#### eQTL overlap with variants reported in GWAS studies of AD

The summary statistics of 4 recent large studies of AD outlined in Table 2 were downloaded from the the GWAS Catalogue ^26^ (version of November 2023). SNPs identified as eQTL in our analysis and reported in the GWAS with a p-value ≤ 0.0001 (GWAS follow up threshold) were identified.

#### pQTL analysis

eQTLs reported in GWAS as associated with AD were tested for association with any of the 741 proteins in the Olink Neurological panel to gain functional insights by understanding the genetic basis of the regulation of proteins involved in the development or progression of AD. A SNP was designated as a pQTL when its association with a protein yielded an FDR-adjusted p-value of ≤ 0.05.

Both eQTL and pQTL analyses were conducted utilizing the *MatrixEQTL* ^27^ R library. *MatrixEQTL* applies a regression model with mRNA/protein levels as the outcome and additive genotypes as independent variables. The regression model was adjusted for covariates, including age, sex, and principal components (PCs) 1 to 6 that explain most of the variance, to account for genetic ancestry admixture.

## RESULTS

### Data Description

GENE-FORECAST and MH-GRID data were combined in the eQTL and pQTL analyses to maximize the sample sizes. A description of the baseline characteristics of the 342 GENE-FOREAST and 133 MH-GRID samples included in the eQTL analysis are outlined in Table 1.

**Table 1:** Four recent large GWAS studies of AD with summary statistics available for download from the GWAS catalogue.

| GWAS Catalogue Study Accession | PubMed ID | Publication Date (Author) | Study Title | Sample Size |
| --- | --- | --- | --- | --- |
| GCST9002715 | 35379992 | 2022-04-04 (Bellenguez et al.) | New insights into the genetic etiology of Alzheimer's disease and related dementias | 111,326 cases<br>677,663 controls |
| GCST013196 | 34493870 | 2021-09-07 (Wightman et al.) | A genome-wide association study with 1,126,563 individuals identifies new risk loci for Alzheimer's disease | 90,338 cases<br>1,036,225 controls |
| GCST007320 | 30617256 | 2019-01-07 (Jansen et al.) | Genome-wide meta-analysis identifies new loci and functional pathways influencing Alzheimer's disease risk | 71,880 cases 383,378 controls |
| GCST005922 | 29777097 | 2018-05-18 (Marioni et al.) | GWAS on family history of Alzheimer's disease | 39,918 cases<br>101,742 controls |

**Table 2:** Baseline characteristics of GENE-FORECAST and MH-GRID samples included in.

| Characteristics | GENE-FORECAST (n = 342) |  | MH-GRID (n = 133) |  |
| --- | --- | --- | --- | --- |
|  | Mean or Count | SD or Proportion | Mean or Count | SD or Proportion |
| Age (years) | 48 | 12 | 45 | 7 |
| Sex |  |  |  |  |
| Female | 235 | 69% | 45 | 34% |
| Male | 107 | 31% | 88 | 66% |
| Systolic Blood Pressure, SBP (mmHg) | 154 | 38 | 120 | 16 |
| Diastolic Blood Pressure, DBP(mmHg) | 76 | 10 | 77 | 11 |
| Body Mass Index, BMI (kg/m2) | 32 | 7 | 31 | 9 |
| Low Density Lipoprotein, LDL (mg/dL) | 112 | 39 | 109 | 32 |
| Fasting Blood Glucose, FBG (mg/dL) | 103 | 35 | 90 | 9 |
| Current Smoker |  |  |  |  |
| No | 399 | 87% | 68 | 51% |
| Yes | 58 | 13% | 62 | 47% |

### Principal component analysis

Because the analysis combined two datasets, principal component analysis (PCA) was undertaken to ensure the two datasets are homogeneous across the transcriptome and proteome data analyzed and the ensuing eQTL and pQTL results are not due to batch effects. The plots in Figure 1 indicate that the two datasets cluster together across the 28 taste perception-related mRNAs and the 741 proteins in the Olink’s neurological panel.

**Figure 1:**
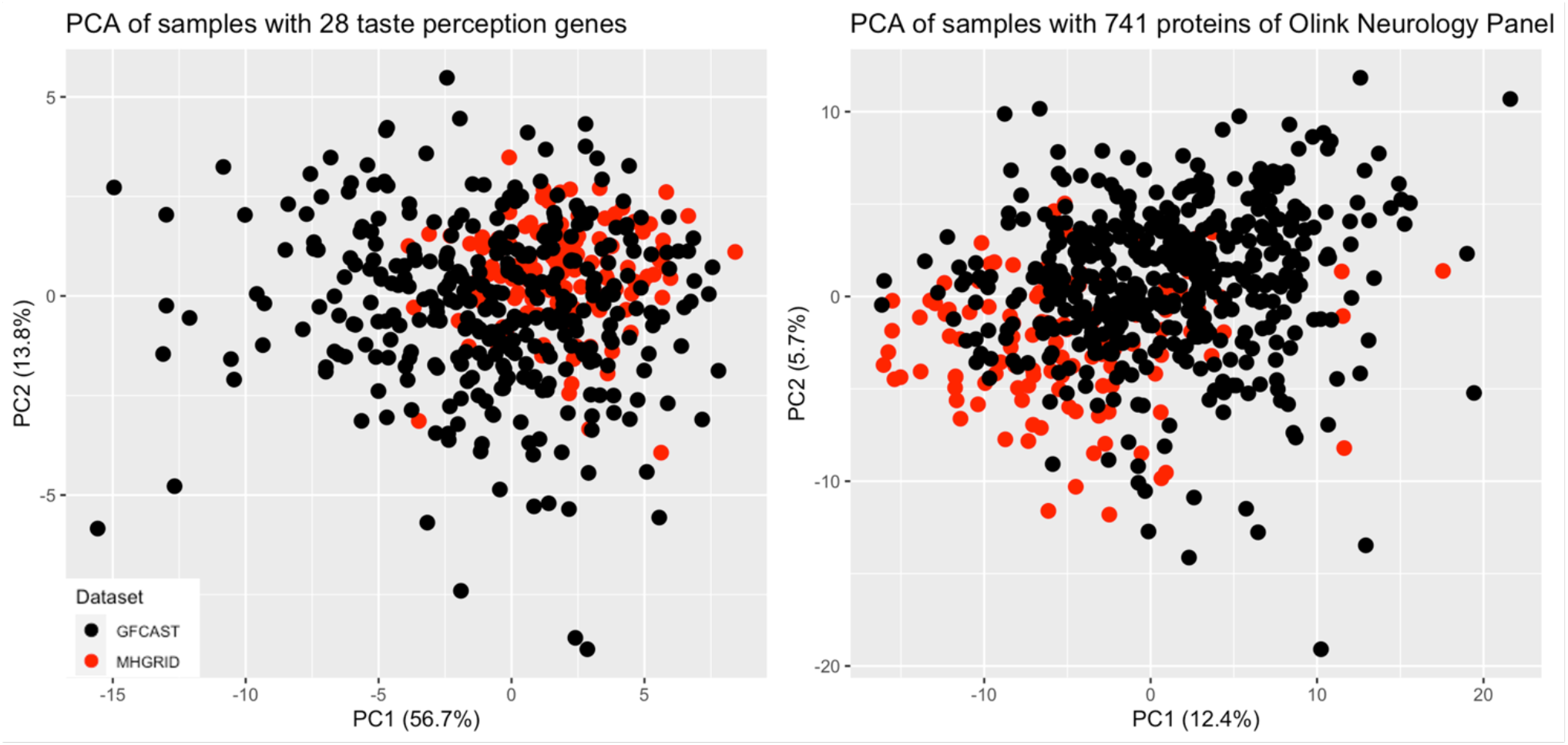
PCA results indicate that the two datasets combined in this analysis cluster well together across PC1 which explains most the variance within the data.

### eQTL analysis and overlap with GWAS findings

A comprehensive total of 3,547 SNP-mRNA associations were statiscally significant after adjustment for multiple testing. These associations comprised of 412 distinct SNPs and encompassed all 28 taste genes. Details of all the eQTL associations and the number of eQTL associated with each gene are outlined respectively in Supplemental Tables T1B and T1C.

A set of 11 eQTLs associated with 16 mRNAs, in the eQTL analysis, were reported in 17 independent GWAS studies of a dozen traits. The intersection with GWAS, succinctly presented in Table 3, includes three SNPs (rs11771145, rs11762262, rs10228407) consistently associated with AD across multiple studies (Table 4). The comprehensive information of SNP, mRNA and GWAS is provided in Supplemental Tables T1D.

**Table 3:** GWAS studies reporting associations with SNPs revealed as eQTL linked to taste genes in our analysis.

| GWAS Trait/Disease | PubMed IDs | SNPs |
| --- | --- | --- |
| Bitter taste perception (multivariate analysis) | 30223776 | rs10772420, rs10261515 |
| Bitter taste perception | 23966204, 30223776 | rs1031391, rs10772420 |
| Idiopathic intracranial hypertension | 29608535 | rs200288366 |
| Height | 18391951, 25282103, 20881960, 23563607 | rs2187642, rs2856321 |
| Alzheimer's disease | 35379992, 31473137, 34493870 | rs11771145, rs11762262, rs10228407 |
| Alzheimer's disease (late onset) | 24162737, 30617256 | rs11771145, rs11762262, rs10228407 |
| Alzheimer's disease or family history of Alzheimer's disease | 29777097, 30617256 | rs11771145, rs11762262, rs10228407 |
| Body size at age 10 | 32376654 | rs2187642 |
| Tea with sugar liking | 35585065 | rs10772380 |
| Gamma glutamyl transferase levels | 33462484 | rs11978404 |
| Bitter taste perception (phenylthiocarbamide) in obesity with metabolic syndrome | 31005965 | rs13231650 |
| Type 2 diabetes | 35893037 | rs4920461 |

**Table 4:** Effect allele, effect size and p-values reported by the 4 GWAS studies of AD for the 3 SNPs identified as eQTLs associated with TAS2R41 and TAS2R60. Two of the SNPs are not reported by Marioni et al.

| GWAS Study | GWAS stats/info | rs11762262 | rs11771145 | rs10228407 |
| --- | --- | --- | --- | --- |
| **Wightman et al.**<br>**PMID 34493870** | log Odds | -0.07 | -0.05 | 0.06 |
|  | P-Value | 9.80e-08 | 3.81e-07 | 2.71e-08 |
|  | Mapped Gene (location) | EPHA1-AS1 (intron) | EPHA1-AS1 (intron) | EPHA1-AS1 (intron) |
|  | Effect allele reported | T | A | T |
| **Marioni et al.**<br>**PMID 29777097** | log Odds | Not reported | Not reported | 0.05 |
|  | P-Value | Not reported | Not reported | 1.28e-09 |
|  | Mapped Gene (location) | Not reported | Not reported | EPHA1-AS1 (intron) |
|  | Effect allele reported | Not reported | Not reported | T |
| **Bellenguez et al.**<br>**PMID 35379992** | Odds-Ratio | 0.94 | 0.94 | 1.05 |
|  | P-Value | 6.35e-10 | 1.29e-12 | 1.24e-10 |
|  | Mapped Gene (location) | EPHA1-AS1 (intron) | EPHA1-AS1 (intron) | EPHA1-AS1 (intron) |

|  | Effect allele reported | T | A | T |
| --- | --- | --- | --- | --- |
| **Jansen et al. PMID 30617256** | log Odds | -0.016 | -0.011 | -0.01 |
|  | P-Value | 5.22e-09 | 8.84e-07 | 9.33e-06 |
|  | Mapped Gene (location) | EPHA1-AS1 (intron) | EPHA1-AS1 (intron) | EPHA1-AS1 (intron) |
|  | Effect allele reported | T | A | G |

In our eQTL analysis all three SNPs exhibit significant association with a down-regulation of TAS2R41 and two of them (rs11771145, rs10228407) are additionally associated with the down-regulation of TAS2R60, reported in Table 5 and Figures 2. The three SNPs have a common frequency in the analysis datasets; they are all upstream of TAS2R41 and TAS2R60 and are not in significant linkage disequilibrium (LD), in the analysis dataset. The LD values measured as r^2^ are respectively 0.58 (between rs11771145 and rs10228407), 0.06 (between and rs11771145 and rs10228407) and 0.10 (between and rs11771145 and rs11762262).

**Table 5:** eQTL analysis results for the 3 SNPs associated with AD in multiple GWAS studies.

| SNP/eQTL | Chr. | Position (GRCh38) | Location | mRNA | Beta | P-Value (raw) | P-Value (adjusted) | Allele tested (Minor Allele) | MAF |
| --- | --- | --- | --- | --- | --- | --- | --- | --- | --- |
| rs11762262 | 7 | 143410783 | 67.1 kb upstream | TAS2R41 | -0.07 | 9.45e-04 | 1.99e-02 | A | 0.21 |
| rs11771145 | 7 | 143413669 | 64.2 kb upstream | TAS2R41 | -0.15 | 2.58e-21 | 3.39e-19 | G | 0.43 |
| rs11771145 | 7 | 143413669 | 29.8 kb upstream | TAS2R60 | -0.33 | 3.26e-27 | 5.77e-25 | G | 0.43 |
| rs10228407 | 7 | 143430678 | 12.8 kb upstream | TAS2R60 | -0.31 | 7.40e-24 | 1.05e-21 | A | 0.46 |
| rs10228407 | 7 | 143430678 | 47.2 kb upstream | TAS2R41 | -0.15 | 5.11e-20 | 6.40E-18 | A | 0.46 |

**Figure 2:**
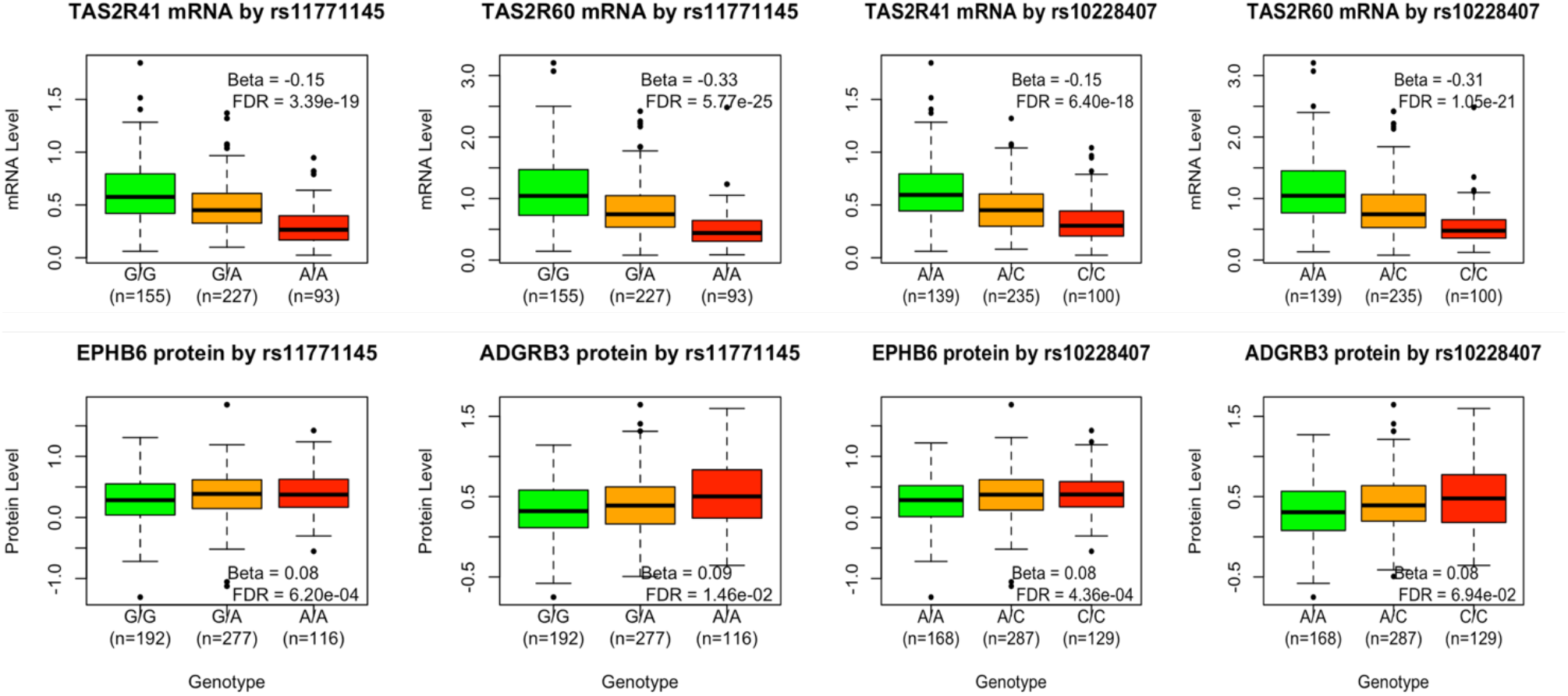
mRNA expression and protein level by rs11771145 and rs10228407 genotypes. The beta and FDR adjusted p-value of each association is provided in the legend.

### pQTL analysis of the 3 SNPs reported in GWAS of AD

The association between all 741 proteins in the Olink’s neurological panel and the 3 SNPs was evaluated. A set of three proteins (EPHB6, ARHGEF5, KEL) are encoded by genes in the vicinity (cis) of the three SNPs and the remaining 738 proteins are encoded by genes further away or in another chromosome (trans). The pQTL analysis revealed that rs11771145 was associated with an upregulation of one cis protein (EPHB6, beta=0.8, p-value=0.0002, FDR adjusted p-value=0.0006) and one trans protein (ADGRB3, 0.09, p-value=0.00002, FDR adjusted p-value=0.014) whilst rs10228407 was associated with an upregulation of cis EPHB6 (beta=0.08 and p-value=0.0001, FDR adjusted p-value=0.0004) and associated with trans ADGRB3 (beta=0.08 and p-value=0.00009, FDR adjusted p-value=0.069). These results are depicted graphically in Figure 2.

## DISCUSSION

The detection of bitter taste receptors such as *TAS2R41* and *TAS2R60* beyond the oral cavity has substantially broadened the scope of research revealing their multifaceted biological functions ^28^. Originally evolved to discern noxious compounds, these receptors are now recognized as integral contributors to immunological defense mechanisms ^29^. Notably, their localization within the cerebral cortex and choroid plexus represents a seminal expansion of their previously ascribed functional roles, implying a plausible engagement in metabolic and immune processes within the brain ^30^.

### Neurological Implications

Recent investigations have unveiled the prospect that bitter taste receptors may exert their influence on neurological pathways implicated in neurodegenerative conditions, such as AD ^31^. Traditionally acknowledged as chemical sentinels and more recently recognized as immune modulators ^32^, these receptors offer novel insights into the mechanisms governing the pathophysiology of neurodegenerative diseases ^33^. Their involvement in the immune and metabolic regulation within the brain not only broadens our understanding but also prompts the consideration of these receptors as potential therapeutic targets or early-stage biomarkers for neurodegenerative disorders ^34^.

### Genetic Associations Reported

Our research has brought to light noteworthy genetic associations linked to the *TAS2R41* and *TAS2R60* genes-associations extending beyond taste perception to intersect with AD. Specifically, the SNPs rs11771145 and rs10228407 located upstream of these genes, emerge as potential critical influencers of their expression. While the primary function of *TAS2R41* and *TAS2R60* is rooted in gustatory detection ^35^, the association between these SNPs and AD is consistently reported in large GWAS investigations which beckons further explorations into the broader implications of genetic variants. Studies suggest that TAS2Rs might have broader implications for health and disease, including their expression in the brain and potential roles in neuronal signaling and disease pathology ^36^. The association of TAS2Rs SNPs and their potential modulation during AD could shed light on the mechanisms of taste alterations and offer new perspectives on how sensory changes are linked to neurodegenerative processes.

### Immunomodulation, Neuroinflammation and Alzheimer’s disease

Previous investigations have highlighted the role of the family of TAS2R genes in modulating immune cell activity, specifically in the regulation of antimicrobial peptides and inflammatory responses ^37^. If these receptors extend a similar influence within the central nervous system, they could potentially modify microglial activity, thereby impacting neuroinflammatory pathways. The crucial function of bitter taste receptors in attenuating neuroinflammatory responses under normal physiological conditions is now well-established ^38^. Conversely, a reduction in the expression of pivotal components within the taste receptor signaling pathway may escalate oxidative stress and activate the inflammasome, ultimately leading to neuroinflammation ^39^.

Neuroinflammation, a shared characteristic among various neurodegenerative diseases, including AD, is now acknowledged as central to their pathophysiology ^40^. The involvement of TAS2R bitter taste receptors in inflammatory responses underscores their significance beyond the realm of taste perception ^41^. Pioneering work by Du et al. in 2018 unveiled that the loss of α-gustducin, a G-protein subunit integral to taste signal transduction, resulted in inflammatory responses and tissue damage ^42^. This underscores the pro-inflammatory potential associated with disruptions in taste receptor signaling, specifically through the NF-κB signaling pathway and the NLRP3 inflammasome ^43^.

### Genetic Variations and Alzheimer’s Pathology

The identification of significant SNP-Gene associations across all 28 taste genes, particularly those leading to the down-regulation of *TAS2R41* and *TAS2R60*, presents a notable link to Alzheimer’s disease within the African American cohort studied. The location of rs117771145 and rs10228407 within the regulatory regions of *TAS2R41* and *TAS2R60*, coupled with their association with AD, in GWAS, fortifies the proposition that bitter taste receptors may be involved in the pathological processes of AD. Variations in expression mediated by these SNPs have the potential to modulate receptor activation thresholds, thereby influencing cellular processes pivotal to neuroprotection, management of neuroinflammatory responses, and clearance of amyloid-beta ^44^. Consequently, such variations may exacerbate neuronal susceptibility to the characteristic pathologies of AD.

### Proteomic Insights

To gain further functional insights, we conducted a comprehensive pQTL analysis targeting three SNPs, revealed as eQTL associated with *TAS2R45* and *TAS2R60* in our analysis, and previously identified in GWAS of AD ^45^. This exploration unveiled compelling associations, elucidating proteins subject to modulation by these specific SNPs. Particularly noteworthy was the observation that rs11771145 exhibited a pronounced upregulation of EPHB6 (Ephrin type-B receptor 6), a protein encoded by a gene within the cis-region of the SNP involved in many developmental processes including neuronal development, angiogenesis, and cell migration ^46^. This observation posits a compelling inference of direct regulatory influence. In the context of AD, EPHB6 assumes prominence owing to its extensive relavance in synaptic plasticity ^47^, neuroprotection, and neuroinflammation ^48,49^. Its regulatory role in fundamental processes positions EphB receptors as potential modulators of AD pathophysiology ^50^.

Moreover, rs11771145 manifested an intriguing association with a protein, encoded by a gene, *ADGRB3* (Adhesion G Protein-Coupled Receptor B3, also known as BAI3), located on a another chromosome. This finding implies the potential for long-range interactions that exert influence over protein expression levels. These associations collectively suggest that the implicated SNPs not only correlate with alterations in gene expression but also wield downstream effects on protein levels. ADGRB3 has been implicated in synaptic regulation and may influence neural circuit formation and plasticity. Alterations in ADGRB3 expression or function have been linked to various neurological conditions, suggesting that it plays a role in maintaining normal cognitive and neural functions ^51^. The brain-specific angiogenesis inhibitor 1 (BAI1), also known as Adhesion G protein-coupled receptor B1 (ADGRB1), emerges as a pivotal regulator of synaptic plasticity ^52^, particularly in the hippocampus. Its involvement in learning and memory processes underscores its significance ^53^. Furthermore, ADGRB1 has been implicated in neuroprotection, mitigating toxin-induced neuronal cell death, and has known associations with dopaminergic neuronal loss in Parkinson’s disease ^54,55^. Concurrently, ADGRB3, enriched in post-synaptic density and cerebellar Purkinje cells ^56^, orchestrates synaptic connections, particularly within the cerebellum ^57^. Genetic variations in ADGRB3, encompassing SNPs and gene amplifications, have been linked to familial schizophrenia and other psychiatric conditions, including bipolar disorder ^58,59^.

The association of the identified SNPs with upregulation of EPHB6 and ADGRB3 proteins suggests potential pathways through which genetic variations may contribute to the unique progression of AD in African Americans. As noted EPHB6 and ADGRB3 are involved in neuronal function and development, and their dysregulation could have significant implications for neurodegeneration. In both cases, the specific mechanisms by which *TAS2R41* and *TAS2R60* influence the expression or function of EPHB6 and ADGRB3 in the context of Alzheimer’s disease remain to be fully elucidated. It is possible that changes in TAS2R expression alter signaling cascades or cellular environments in ways that impact these proteins, which in turn could affect neuronal health and function. Further research is needed to understand these relationships and their implications for AD and sensory health.

### Integrated Perspective and Potential Therapeutic Implications

This integrative perspective, elucidating the interplay between SNPs, taste receptor genes and AD through pQTL analysis, offers profound insights into the intricacies of AD’s etiology. Beyond merely illuminating potential mechanistic pathways, it underscores the multifaceted nature of neurodegenerative diseases. This holistic approach encourages nuanced research strategies for the refinement of therapeutic interventions and the development of preventive strategies.

Furthermore, the neuroprotective and anti-inflammatory attributes associated with bitter compounds, known to interact with TAS2Rs, substantiate the hypothesis that these receptors could potentially modulate the pathophysiological conditions of AD ^60^. Compounds like flavonoids and polyphenols, which engage with TAS2Rs, have exhibited promise in enhancing cognitive function and mitigating markers of neurodegeneration in disease models ^61^. The exploration of TAS2Rs in orchestrating the therapeutic effects of these compounds represents an intriguing avenue for further investigation ^62^.

### Limitations

Our study, elucidating novel insights into the genetic foundations of taste perception genes and their correlation with AD in an African American cohort, is constrained by several limitations. AD was not assessed in this cohort and it was hence not possible to directly link the mRNA and protein changes to AD.

Whole-blood transcriptome may imperfectly reflect brain gene expression, the primary site of AD pathology.

An additional limitation is our reliance on GWAS data predominantly derived from European populations. This underrepresentation of African American and other non-European populations in genetic research can limit the generalizability of our findings. The genetic architecture of AD may vary across different ancestries, and findings from European cohorts may not fully capture the genetic risk factors pertinent to populations of African ancestry. However, the main variants we reported are all common, suggesting they are not ancestry-specific and that our findings might be transferable across populations. Future studies should prioritize the inclusion of diverse populations to enhance the applicability of genetic research across different ethnic groups and to address health disparities in the understanding and treatment of AD.

The cross-sectional design hampers capturing the dynamic nature of gene expression changes over the disease course. Unaccounted environmental and lifestyle factors may confound genetic associations. Lack of direct correlation between genetic variations and clinical manifestations of AD, absence of detailed pathway analysis for SNP-gene and SNP-protein associations, and omission of microRNA exploration further limit our study. Notably, functional consequences of identified expression and protein QTLs lack experimental validation. Future research necessitates in vitro studies to discern genetic variant impact on gene expression and protein levels and function. Subsequent in vivo studies are vital for establishing their role in disease processes, fostering a comprehensive understanding of these associations in AD context.

## Conclusions

Our investigation has unveiled novel insights into the genetic determinants implicated in AD, underscoring the potential important role of bitter taste receptor genes in its pathophysiological mechanisms. The identification of SNPs exhibiting robust associations with both taste perception genes and AD provides evidence for the plausible involvement of these genes in the intricate pathways of neurodegenerative diseases ^63^. These findings posit genetic variations influencing taste receptor expression as potential contributors to brain health, positioning these receptors as plausible biomarkers for AD ^64^.

The implications of our study for unraveling the genetic architecture of Alzheimer’s disease have valuable significance, offering prospects for the development of targeted interventions aimed at modulating the pathways identified. Our findings also raise new questions and avenues for future research. Subsequent investigations should prioritize the validation of these associations within broader and more genetically diverse cohorts to ascertain the generalizability of our observations. Furthermore, longitudinal studies are warranted to elucidate the causal relationships between these genetic variations, gene expression regulation, and the progression of AD.

Our study highlights a significant step towards understanding the complex interplay of genetics, sensory function, and neurodegeneration in Alzheimer’s disease within an African American cohort. Critical to the translation of genetic insights into therapeutic strategies, future research endeavors should delve into the direct impact of these SNPs on the neural circuitry and cognitive functions affected by AD. Such investigations will be instrumental in advancing our understanding and, consequently, facilitating the development of precise therapeutic interventions.

## ETHICS APPROVAL AND CONSENT TO PARTICPATE

This study was approved by National Institutes of Health Institutional Review Board (IRB). The study was conducted in accordance with the local legislation and institutional requirements. The participants provided their written informed consent to participate.

## CONSENT FOR PUBLICATION

Not applicable.

## AVAILABILITY OF DATA AND MATERIAL

Additional information can be found in the Supplementary Material of this article. The datasets presented in this article cannot be publicly shared due to privacy restrictions. Requests to access the datasets should be directed to the corresponding author.

## AUTHOR’S CONTRIBUTIONS

AG and PVJ designed the analysis. GG processed and conducted quality controls of the transcriptome, proteome and phenotype data. AG conducted the statistical analyses. AG, PVJ and MA interpreted the results. AG, PVJ, MA, GG and AD drafted and edited the manuscript. All authors reviewed and approved the final version of the manuscript.

## ACKNOWLEDGEMENTS

PVJ is supported by National Institute of Alcohol Abuse and Alcoholism under award number, Z01AA000135, the National Institute of Nursing Research and the Rockefeller University Heilbrunn Nurse Scholar Award. PVJ is supported by the Office of Workforce Diversity, and the Office of Workforce Diversity, National Institutes of Health Distinguished Scholar Program.

We are grateful to Dr Gary H. Gibbons, the initial PI of the GENE-FORECAST study.

## ABBREVIATIONS

eQTL: Expression-Quantitative Trait Loci
pQTL: Protein-Quantitative Trait Loci
GWAS: Genome-Wide Association Study
SNP: Single-Nucleotide Polymorphisms
AD: Alzheimer’s Disease
GENE-FORECAST: GENomics, Environmental FactORs and the Social DEterminants of Cardiovascular Disease in African-Americans Study
MH-GRID: Minority Health Genomics and Translational Research Bio-Repository Database
MSM: Morehouse School of Medicine
PCA: Principal Component Analysis
LD: Linkage Disequilibrium
SBP: Systolic Blood Pressure
DBP: Diastolic Blood Pressure

